# Adaptation during the transition from *Ophiocordyceps* entomopathogen to insect associate is accompanied by gene loss and intensified selection

**DOI:** 10.1101/2024.03.04.583259

**Authors:** Chris M. Ward, Cristobal A. Onetto, Anthony R. Borneman

## Abstract

Fungal and bacterial symbiosis is an important adaptation that has occurred within many insect species, which usually results in the relaxation of selection across the symbiont genome. However, the evolutionary pressures and genomic consequences associated with this transition are not well understood. Pathogenic fungi of the genus *Ophiocordyceps* have undergone multiple, independent transitions from pathogen to associate, infecting soft-scale insects trans-generationally without killing them. To gain an understanding of the genomic adaptations underlying this transition, long-read sequencing was utilized to assemble the genomes of both *Parthenolecanium corni* and its *Ophiocordyceps* associate from a single insect. A highly contiguous haploid assembly was obtained for *Part. corni*, representing the first assembly from a single Coccoidea insect, in which 97% of its 227.8 Mb genome was contained within 24 contigs. Metagenomic-based binning produced a chromosome-level genome for *Part. corni*’s *Ophiocordyceps* associate. The associate genome contained 524 gene loss events compared to free-living pathogenic *Ophiocordyceps* relatives, with predicted roles in hyphal growth, cell wall integrity, metabolism, gene regulation and toxin production. Contrasting patterns of selection were observed between the nuclear and mitochondrial genomes specific to the associate lineage. Intensified selection was most frequently observed across nuclear orthologs, while selection on mitochondrial genes was found to be relaxed. Furthermore, scans for diversifying selection identified associate specific selection within three adjacent enzymes catalyzing acetoacetate’s metabolism to acetyl-COA. This work provides insight into the adaptive landscape during the transition to an associate life history, along with a base for future research into the genomic mechanisms underpinning the evolution of *Ophiocordyceps*.

## Introduction

Microbes interact with higher organisms through diverse and specific mechanisms (Iturbe-Ormaetxe, et al. 2011; Fan, et al. 2015; Flórez, et al. 2017; Kobmoo, et al. 2018; Trivedi, et al. 2020) adapting through both antagonistic and mutualistic strategies. Mutualism between microbes and associated taxa exists whereby fitness benefits are conferred to both organisms by mutual and generally symbiotic, relationships. In insects, an association with specific prokaryotic and fungal species has been shown to provide mutual benefit to the host through increasing nutrient availability (Noda and Koizumi 2003; Vogel and Moran 2011) or detoxification of noxious compounds (Welte, et al. 2016). As the mutualistic relationships can insulate the associate from direct environmental influences, which can drastically change selection pressures that are applied to the symbiont genome and allow for previously constrained genes to escape selective pressures (Wernegreen 2011; Wertheim, et al. 2015). This is most dramatically observed when symbionts live within the body of the insect host or even within the insect cells (Wertheim, et al. 2015) and are inherited vertically, drastically decreasing the effective population size, while removing the need for sexual reproduction (Mira and Moran 2002). The loss of sexual recombination during the switch from a free-living to an asexually-inherited obligate lifestyle results in the accumulation of deleterious mutations via Muller’s Rachet (Muller 1964). For example, facultative and obligate γ-proteobacteria endosymbionts of insects show clear signals for relaxed selection when compared to their free-living relatives (Wertheim, et al. 2015). In addition to selective relaxation, gene loss and genomic minimization are a common feature of the transition to an obligate lifestyle, with the loss of genes that encode now superfluous metabolic pathways (Osvatic, et al. 2023), in addition to those involved in DNA regulation or repair (Shigenobu, et al. 2000). Accumulation of deleterious mutations can also drive pseudogenization, whereby genes lose functionality while being retained in the genome (van der Burgt, et al. 2014; Esfeld, et al. 2018).

Fungal symbiosis has been described throughout insects, with many obligate fungal species providing benefits to their host (Dowd 1992; Noda and Koizumi 2003; Douglas 2015; Stefanini 2018). *Ophiocordyceps* are entomopathogenic and nematopathogenic fungi that have evolved highly complex life histories to target, control and kill their host (Kobmoo, et al. 2018). Many species are highly host-specific, with species complexes evolving via unique mechanisms to infect their host (Evans, et al. 2011; Beckerson, et al. 2023). However, in soft-scale insects (Hemiptera:Coccomorpha) *Ophiocordyceps* have undergone multiple independent transitions from a pathogenic to associate lifestyle, where they are inherited vertically from female insects directly to their eggs (Szklarzewicz, et al. 2021). *Ophiocordyceps* associates have been demonstrated to live within the lipid rich storage tissues of their host and upon female maturation and egg production, migrate to the ovum to infect the eggs as they are forming (Szklarzewicz, et al. 2021).

In this study long-read sequencing was used to produce a highly contiguous genome from a single *Parthenolecanium corni* (Coccomorpha:Coccidae) individual along with its *Ophiocordyceps* obligate. Genomic analysis of the *Part. corni* genome identified multiple structural rearrangements and a decrease in the frequency of the conserved Mariner repeat family. Phylogenetic reconstruction of the *Ophiocordyceps* obligate confirmed its closest ancestor as *Hirsutella rhossiliensis* and identified widespread gene loss and signals for intensified selection within the nuclear genome of the fungal associate, while in contrast, the mitochondrial genome exhibited signals for relaxed selection. This work provides the first genome assembly of an obligate *Ophiocordyceps* and suggests widespread intensified selection coupled with diversifying selection on enzymes in the acetoacetate metabolism as key players in the transition from pathogenic to associate lifestyles in this system.

## Methods

### Sample collection, DNA extraction and sequencing

A single *Parthenolecanium* corni individual was collected from a grapevine in the Langhorne Creek wine region of South Australia. The *Part. corni* individual was inspected under a stereomicroscope and washed using 95% ethanol and brushed three times to remove potential environmental contaminants. DNA was then extracted via gentle homogenization of the insect tissue using a glass rod in lysis buffer followed by 24-hour proteinase-K digestion, RNAse A incubation, a modified phenol-chlorophorm extraction (Ward, et al. 2021) and glycogen/isopropanol precipitation at 4°C. DNA molecular weight and quality was determined using a TapeStation (Agilent) at the Australian Genome Research Foundation. A total of 222 ng of DNA was obtained and used to prepare a nanopore library using the SQK-LSK114 kit. The library was loaded into a FLO-PRO114M flow-cell and sequenced in a P2-Solo instrument. Basecalling was performed using Guppy v.6.4.6 with the SUP (r10.4.1_e8.2_400bps_sup) model. Duplex paired reads were identified using duplex-tools and re-basecalled using Guppy. After quality filtering (Q > 10) a total sequencing yield of 76.17 Gb (71.98 simplex and 4.19 Gb of duplex reads) was obtained.

### k-mer analysis

Simplex reads were used to carry out k-mer analysis using Canu v2.2 (Koren, et al. 2017) to investigate 22-mer depth distribution and estimate ploidy and genome size. Genome size estimation was carried out by calculating the area under the curve for local maxima present in the 22-mer depth distribution.

### Genome assembly and separation

Metagenome assembly was carried out using Meta-Flye v2.9.3 (Kolmogorov, et al. 2020) using the combined simplex and duplex reads under default settings. Combined simplex and duplex reads were then mapped to the metagenome using Minimap2 v2.26 (Li 2018) with the map-ont preset. Mapped reads and the metagenome were then passed to MetaBat2 v2.15 (Kang, et al. 2019) for metagenome binning. A combined approach was used to determine if binned subgenomes should be combined. First; binned sub-genome average read depth and variance were compared to the 22-mer depth distribution, second; contig GC content distribution was compared between bins, third; a custom database containing all insect, fungal, and bacterial proteins on NCBI was constructed and contigs were queried using Kaiju v1.10.0 (Menzel, et al. 2016) and the BUSCO v4.1.2 (Seppey, et al. 2019) Insecta and Fungi datasets. NgsReports v2.4 (Ward, To, et al. 2020) was then used to visualize and investigate BUSCO outputs to identify differences in completeness between organisms.

Simplex and duplex fastq files were then mapped separately to the complete metagenome with Minimap2 v2.26 (Li 2018) using the map-ont preset without allowing for secondary mapping. Reads with a MAPQ>=20 were extracted from their respective fastq files and separated into subgenome fraction (ie. *Parthenolecanium* or *Ophiocordyceps* subgenome) using seqtk v75 (https://github.com/lh3/seqtk). The *Parthenolecanium* and *Ophiocordyceps* derived reads were then assembled separately using Verkko v2.0 (Rautiainen, et al. 2023) by providing duplex reads as the ‘high accuracy’ and simplex reads as the ‘low accuracy’ long-read arguments. The two Verkko assemblies were then combined along with Meta-Flye contigs that were not assigned to *Parthenolecanium* or *Ophiocordyceps*. All simplex and duplex reads were then mapped to the Meta-Flye/Verkko assembly and contig error correction was carried out using medaka (https://github.com/nanoporetech/medaka). Finally, corrected contigs were reassigned into *Parthenolecanium* or *Ophiocordyceps* derived genomes with SeqKit v2.7.0 (Shen, et al. 2016). BUSCO v4.1.2 (Seppey, et al. 2019) was run at each stage of the assembly process using the Fungi and Insecta databases to determine single copy ortholog fraction being recovered and identify potential contamination.

### Gene prediction and annotation of the Coccoidea and *Ophiocordyceps* genomes

Gene prediction was carried out on the *Parthenolecanium corni* assembly and publicly available Coccoidea genomes without gene models (Sup Table 1) using the braker2 pipeline (Brůna, et al. 2021) providing all Arthropod OrthoDB (Kriventseva, et al. 2019) orthogroups and Hemipteran proteins available on NCBI to generate hints for annotation.

The *Ophiocordyceps* genome was annotated using the Funannotate pipeline (10.5281/zenodo.2604804) by passing *Ophiocordyceps* proteins along with the SwissProt and UniProt databases to generate hints. Gene predictions were then functionally annotated for all genomes using the Funannotate (10.5281/zenodo.2604804) annotation function and Interproscan v5.59_91.0 (Jones, et al. 2014). GO terms were generated using the IPR domain to GO term table available through Gene Ontology (https://current.geneontology.org/ontology/external2go/interpro2go). Kegg terms were then generated using Kegg Mapper (Kanehisa and Sato 2020) and maps constructed using KEGG Reconstruct (Kanehisa 2017).

### Analysis of the *Parthenolecanium* genome

Synteny analysis between the *Part. corni* genome and *Planococcus citri* (GCA_950023065.1) was carried out using MCScanX (Wang, et al. 2012) under default settings using contigs >1Mb in total length. Orthogroups were identified between *Part. corni* and publicly available Coccomopha genomes (Sup Table 1) protein annotations using OrthoFinder v2.2.7 (Emms and Kelly 2019) under default settings.

Repeat annotation was carried out by first modeling denovo repeats in the genome using RepeatModeler v2.0.5 (Flynn, et al. 2020) with the RepBase database 28.05 (Jurka, et al. 2005) and RepeatMasker databases (Chen 2004). Repeats were then annotated and masked in the genome using RepeatMasker v4.1.5 (Chen 2004).

Structural variation was characterized within the *Part. corni* genome by mapping the duplex reads to the *Part. corni* sub genome with Minimap2 v2.26 (Li 2018). Mapped reads were then called using Sniffles v2.2 (Sedlazeck, et al. 2018) to identify structural variants and IGV (Thorvaldsdóttir, et al. 2013) was used to manually inspect read mapping around these loci.

### Analysis of the Ophiocordyceps genome

Synteny analysis between the *Ophiocordyceps* genome and *Cordyceps militaris* was carried out using MCscanX (Wang, et al. 2012) under default settings. Orthogroups were identified between *Ophiocordyceps* and Cordyceps genomes (Sup Table 2) protein annotations using OrthoFinder v2.2.7 (Emms and Kelly 2019) under default settings. A species tree was then constructed by combining individual single-copy ortholog gene trees using ASTRAL-III v5.7.1 (Zhang, et al. 2018). Gene loss analysis was carried out by parsing the OrthoFinder orthogroups output to identify genes present in the closest taxa, *H. rhossiliensis*, as identified using phylogenetic analysis. Two different levels of gene loss were then assessed by extracting genes using geaR v1.0 (Ward, Ludington, et al. 2020). First, genes present in all species within the same clade and second those that are present in all assessed *Ophiocordyceps* and *Cordyceps* genomes. GO and KEGG terms of lost genes were determined by first annotating the *H. rhossiliensis* genome using the same methodologies as the *Ophiocordyceps* associate genome above. Annotations were then parsed for their GO and KEGG pathways. Enrichment analysis of GO terms was carried out using the R package clusterProfiler v3.18 (Yu, et al. 2012).

### Selection testing

Single-copy orthologs between the *Ophiocordyceps* associate of *Parthenolecanium corni* (*OAc*), *Ophiocordyceps sinensis, Hirsutella rhossiliensis* and *Hirsutella minnesotensis* were identified using OrthoFinder v2.2.7 (Emms and Kelly 2019) and translation informed codon alignment was carried out with translatorX v1.1 (Abascal, et al. 2010). Gene alignments were then passed to HyPhy v2.5.59 (Pond, et al. 2005) for positive/diversifying selection testing using the BUSTED model (Murrell, et al. 2015). Two diversifying selection hypotheses were tested: first, only the *OAc* branch was tested for diversifying selection for each gene utilizing the species topology. Second, all other internal and terminal branchers excluding the *OAc* terminal branch were tested for diversifying selection. Genes were flagged as under diversifying selection if a statistically significant signal (pvalue ≤ 0.05) was observed in the *OAc* terminal branch without signal (pvalue > 0.05) detected across all other branches.

Intensification/relaxation of selection was calculated using RELAX model (Wertheim, et al. 2015) in HyPhy v2.5.59 (Pond, et al. 2005) utilizing the gene alignments of *OAc, Ophiocordyceps sine, Hirsutella rhossiliensis* and *Hirsutella minnesotensis* single copy orthologs. The *OAc* terminal branch was set to the foreground and evolutionary rate was determined against all other internal and terminal branches.

Duplicated orthogroups were identified from the nucleotide OrthoFinder output. Orthogroups that were found to be duplicated in only *OAc* were tested for diversifying positive selection using the ABSREL model (Smith, et al. 2015) in HyPhy v2.5.59 (Pond, et al. 2005). Each terminal branch was tested for selection and signal for positive diversifying selection was considered if significant signal (p value ≤ 0.05) in either one or both *OAc* paralogues.

Mitochondrial selection was investigated by using the mold codon model in HyPhy v2.5.59 (Pond, et al. 2005) with the ABSREL (Smith, et al. 2015) and RELAX (Wertheim, et al. 2015) models following the same methodology as nuclear genes.

## Results

### Assembly of DNA sourced from a single scale individual recovers contiguous genomes for *Parthenolecanium corni* and a putative *Ophiocordyceps* associate

Sequencing the total genetic content within a single insect allows assembly of bacterial and/or fungal associates to be co-assembled (Ward and Baxter 2017). Due to the small size of soft-scale insects (<4 mm), recovery of sufficient high molecular weight DNA was challenging, however sufficient DNA was recovered for sequencing on a Nanopore P2 flow-cell, which produced 76.1 Gb of long read data (>q10, 71.9 Gb simplex; 4.2 Gb duplex). After removing shorter reads (<500 bp), 62 Gb (N_50_ of 8.05 kb) was retained for metagenomic assembly and which produced a 261.1 Mb assembly distributed across 1331 contigs and which was subsequently binned into 8 metagenome assembled genomes (MAGs) that varied in size from 212 Mb to 220 kb (Figure 1A, Sup Table 3). Combining MAGs based on taxonomic classification of the bins identified a single Insecta/Hemipteran and two Fungal genomes (Figure 1A) that were predicted to belong to the genera *Parthenolecanium* (Hemiptera), *Aureobasidium* (Dothideales) and *Ophiocordyceps* (Hypocreales), respectively (Figure 1A).

**Figure 1.**
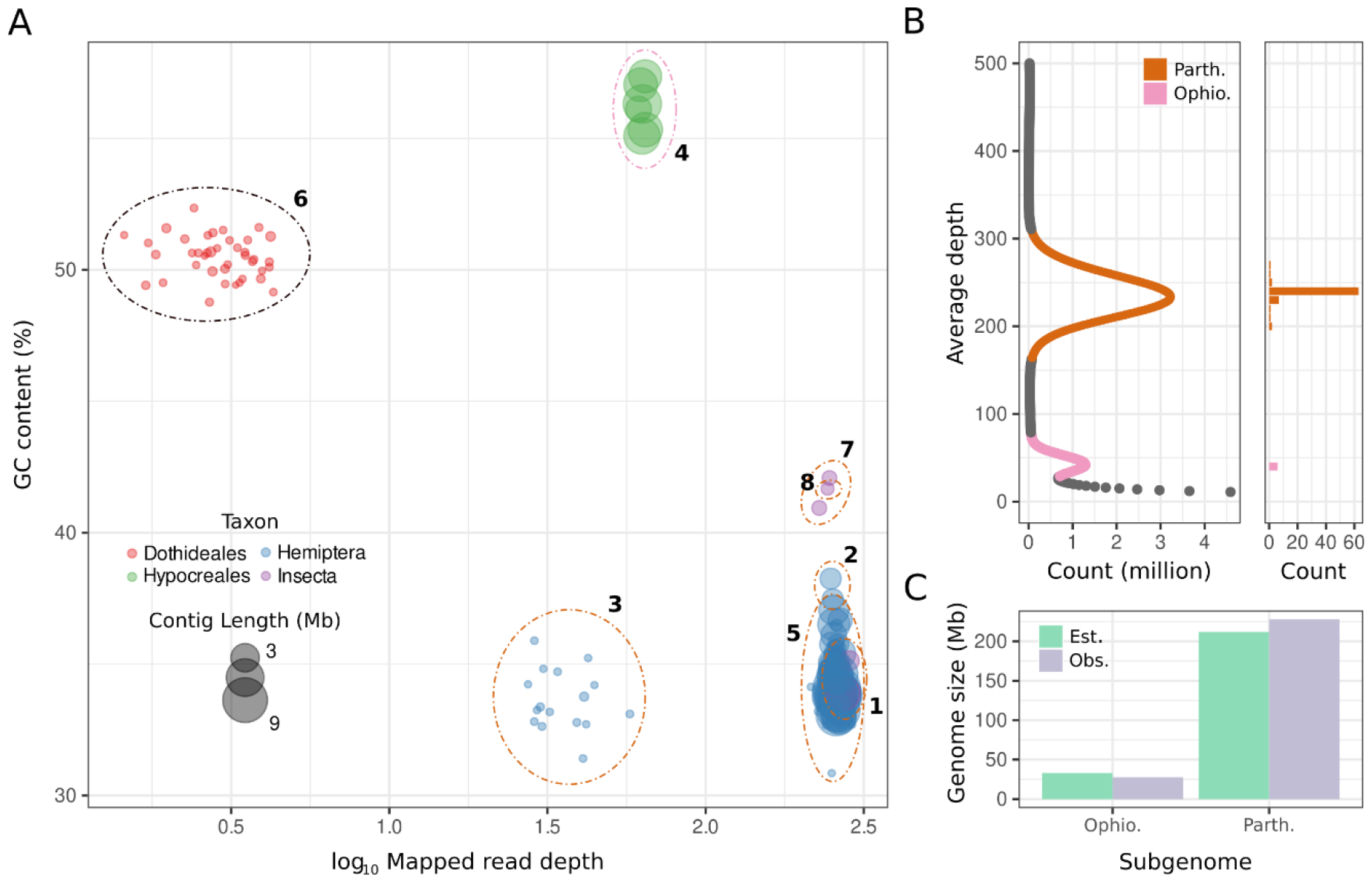
Metagenomic assembly of a single scale insect recovers two contiguous genomes. A) Metagenomic binning of contigs based on contig GC content and read depth. Contigs were then assigned to taxon using flexible protein matches and BUSCO gene content. Contigs are visualized based on contig length in megabases. B) Left: distribution of observed count vs read depth of 21-mers across the genome. Right: number of contigs within each read depth bin for each of the sub genomes. C) Genome size estimation through 21-mer frequency against the observed genome sizes of the *Ophiocordyceps* and *Parthenolecanium* sub-genomes post separation.

Duplex and simplex reads that mapped to the *Ophiocordyceps* and *Part. corni* MAGs were then extracted and used to perform specific assemblies for each organism, producing two highly contiguous genomes that were contained within 6 and 85 contigs, respectively. Assessment of duplex and simplex 22-mer depth revealed two clear peaks in the approximate location of mapped contig read depth distributions (Figure 1B). The 21-mer distribution was then used to determine the estimated genome size of each of the sub-genomes. The difference between recovered and estimated genome size provides insight into over clustering or bin mis-assignment during the assembly and separation stages. Calculating the area under each local maxima estimated the genome size of *Ophiocordyceps* to be 33 Mb and 212 Mb for *Part. corni*, compared to the 27.7 Mb and 227.8 Mb recovered genomes (Figure 1C) suggesting the genomes for both *Part. corni* and its *Ophiocordyceps* associate were well recovered.

### Structural rearrangements and decreased repetitive elements in the *Part. corni* genome

#### Analysis of completeness of the *Part*

*corni* genome revealed similar levels of BUSCO gene presence relative to genome assemblies of other Coccoid species (Sup Figure 1) with 85.6% single copy, 5.5% duplicated (91.1% complete) and 6.8% missing BUSCOs in the *Part. corni* genome. Gene annotation of the *Part. corni* genome identified 14,951 protein coding genes, which is well within the estimated 10,000 - 30,000 present in most insects (Adams, et al. 2000; Holt, et al. 2002; Consortium 2006; Consortium 2010; Ward, et al. 2021).

#### Clustering the *Part*

*corni* genes with other publicly available coccoid genomes identified 23,726 orthogroup clusters containing 86.3% of all genes within the search space, 951 of which represented single copy orthologs present across all genomes. Phylogenetic reconstruction of the *Part. corni* single copy orthologs against the background of mealybug and scale insect coccoid species identified the fellow scale insect *Ericerus pela* as the closest sequenced relative (Figure 2A). Scale insects were found to be in reciprocal monophyly with mealybugs (Figure 2A), agreeing with past literature (Vea and Grimaldi 2016). Synteny analysis of contigs greater than 1Mb in length (n = 24, 97% of total genome size) from the *Part. corni* genome against the chromosome level *Planococcus citri* (mealybug) genome identified almost no macro-synteny at the chromosome level (Figure 2B). To determine if this was due to misassembly in the *Part. Corni* genome, duplex ONT reads were mapped against the *Part. Corni* assembly. One hundred and sixty-nine putative structural variants (SVs) were identified within the *Part. Corni* genome and manual investigation of read alignments surrounding these loci confirmed 13 heterozygous InDels. No evidence for widespread mis-assembly was observed with called major SVs (e.g. inversions and translocations) being caused by mapping errors (Sup Figure 2).

**Figure 2.**
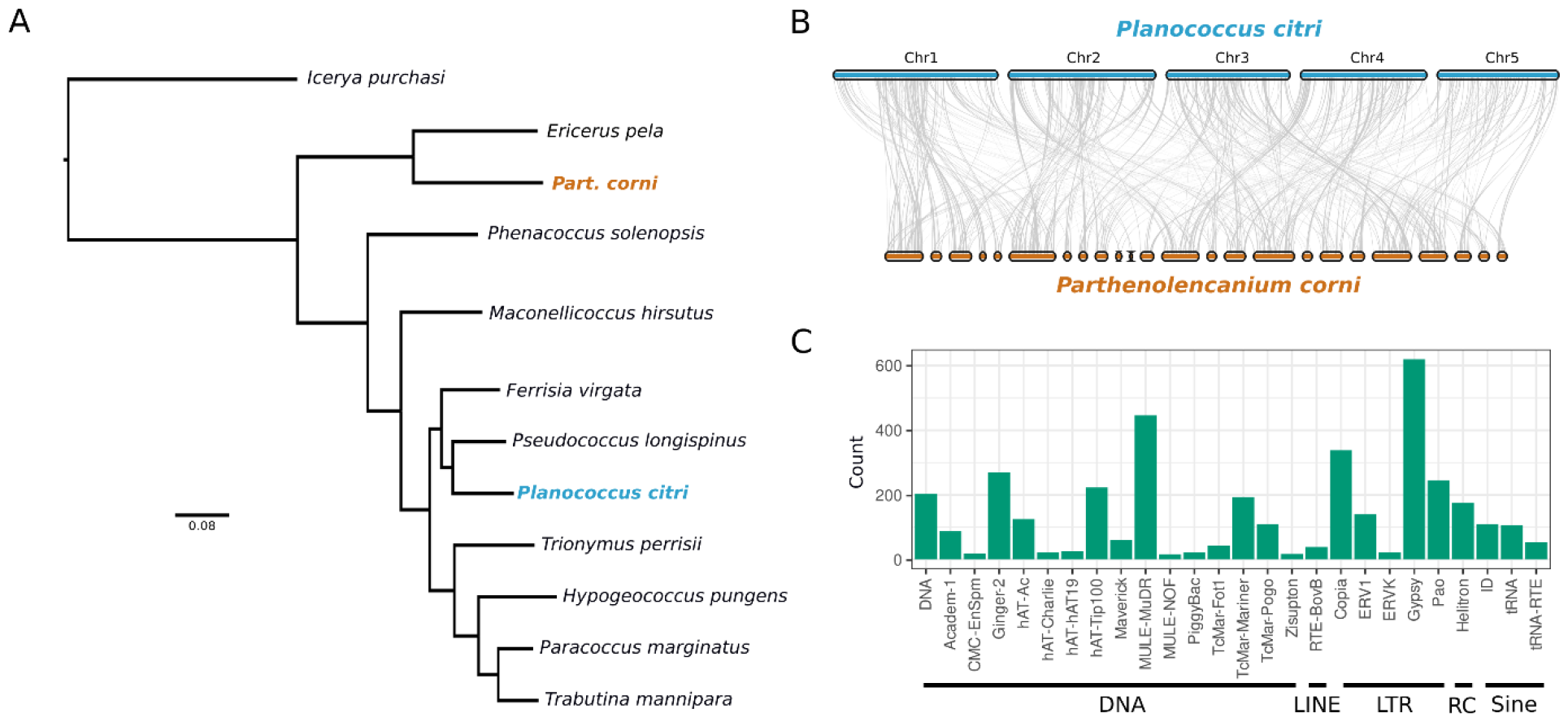
Genome analysis of *Parthenolecanium corni*. A) Phylogenetic reconstruction of Coccoid single copy orthologs. Branch lengths are in coalescent units. B) Comparative synteny analysis of *Planococcus citri* chromosomes (blue) and contigs longer than 1Mb (n = 24) *Parthenolecanium corni* contigs (orange) which made up 97% of the total *Part. corni* genome. C) Repeat content analysis of the *Part*. corni genome. Repeat classes are annotated along the y-axis.

#### Repeat modeling of the *Part*

*corni* genome identified 1,881 DNA, 38 LINE, 268 SINE and 1,364 LTR repetitive elements (Figure 2C), which represents far lower repeat content than the estimates performed on either its nearest neighbor, *E. pela* (Yang, et al. 2019) or the model Hemipteran, *Acyrthosiphon pisum* (Consortium 2010). For example, Mariner repeats are present throughout Hemiptera (Filée, et al. 2015; Bouallègue, et al. 2017; Yang, et al. 2019) and while 812 were observed in the *E. pela* repeat annotation (Yang, et al. 2019), *Part. corni* was found to only have 193.

### The *Ophiocordyceps* associate of *Part. corni* has undergone gene loss in conserved orthologs

*Ophiocordyceps* and *Hirsutella* are synonymous genera, with *Hirsutella* being used to refer to the anamorphic, asexual life stage of this fungus (Simmons, et al. 2015), rather than the two genera representing phylogenetically distinct taxa. Phylogenetic reconstruction of the *Ophiocordyceps* associate of *Part. corni* (referred to as OAc) and publicly available *Ophiocordyceps* and *Cordyceps* genomes identified *Hirsutella rhossiliensis* as the closest sequenced relative to OAc, with high posterior support (Figure 3A). *OAc, H. rhossiliensis, O. sinensis* and *H. minnesotensis* formed a monophyletic clade separate from all other *Ophiocordyceps* genomes (Figure 3A; Clade 1), suggesting that this lineage shared a most recent common ancestor that was separate from other *Ophiocordyceps* species.

**Figure 3.**
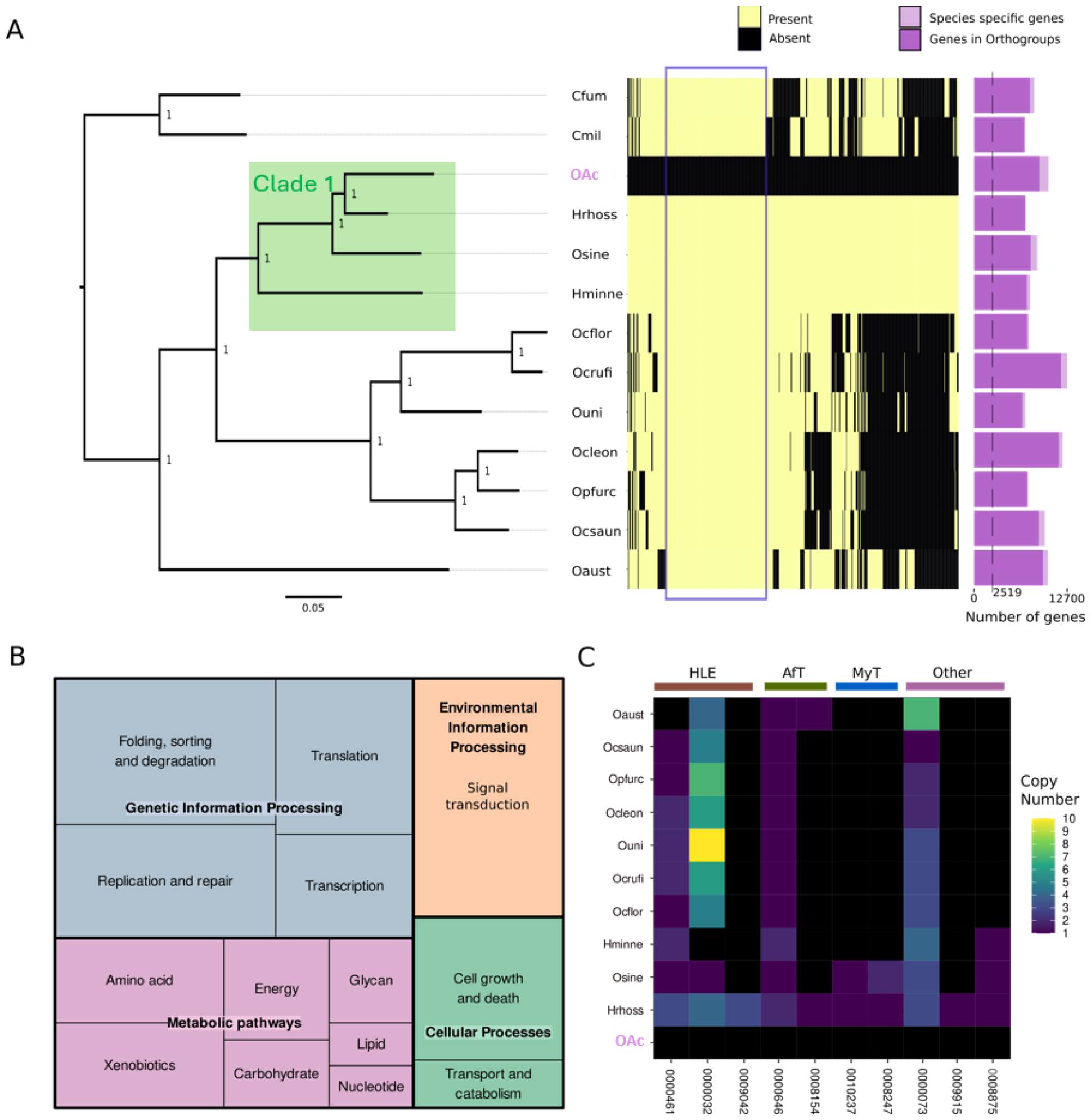
Widespread gene loss of conserved genes in OAc. A) Phylogenetic reconstruction of single copy orthologs of Cordyceps and Ophiocordyceps genomes (Sup Table 2). Gene loss in OAc for orthogroups (x-axis) that were present in the closest phylogenetic neighbors *H. rhossiliensis, O. sinensis* and *H. minnesotensis*. Cfum = *Cordyceps fumosorosea*; Cmil = *Cordyceps militaris*; Hrhoss = *Hirsutella rhossiliensis*; Osine = *Ophiocordyceps sinensis*; Hminne = *Hirsutella minnesotensis*; Ocflor = *Ophiocordyceps camponoti-floridani*; Ocleon = *Ophiocordyceps camponoti-leonardi*; Ocsaun = *Ophiocordyceps camponoti-saundersi*; Oaust = *Ophiocordyceps australis*; Opfur = *Ophiocordyceps polyrhachis-furcata*; Ocrufi = *Ophiocordyceps camponoti-rufipedis*; Ouni = *Ophiocordyceps unilateralis*. Purple box highlights orthogroups where all species contained at least one ortholog that were absent from OAc B) Kegg pathway breakdown of lost genes that are fixed with at least one copy in all entomopathogenic fungi in the tree. C) Loss of toxin related genes in OAc. Heat Labile Enterotoxin (HLE) (OG0000032, OG0000461, OG0009042), Aflatoxin (AFt) (OG0000646, OG0008154), Mycotoxin (MyT) (OG0008247, OG0010237), and Other toxins (OG000073, OG0009915, OG0008875) related gene copy number are shown in species where the gene was observed.

Analysis of the completeness of each genome revealed similar levels of BUSCO gene presence across all of the sequenced *Ophiocordyceps* species (Sup Figure 3) with 96% single copy, 0.9% duplicated and 2.7% missing BUSCOs identified in the OAc genome. The proportion of genes assigned to orthogroups was also similar across the sequenced *Ophiocordyceps*, with no species having a significant number of species-specific genes that were not able to be grouped (Figure 3A). Of the 7357 predicted gene models in OAc, 7247 were successfully assigned to an orthogroup with at least one other *Ophiocordyceps* or *Cordyceps* gene model and the single copy orthogroups present in all species totaled 2319.

Gene gain and loss are essential adaptive strategies employed by many fungi to respond to novel environmental stressors (Osvatic, et al. 2023). To investigate gene loss in the OAc lineage, 524 orthologs were identified that were present in Clade 1 (Figure 3A) but absent in the OAc annotation (Figure 3A). Of these 524 genes, 159 were present in all other taxa throughout the phylogeny (Figure 3B), suggesting that they are likely to play important roles in Hypocreales biology. Genes present in all other taxa are therefore referred to as ‘conserved’ throughout the remainder of the text. OAc exhibited an increased rate of conserved gene loss relative to all other *Ophiocordyceps* species except *O. sinensis* (Sup Figure 4). Functional annotation of the *H. rhossiliensis* orthologs that were absent in OAc revealed roles in amino-acid and xenobiotic metabolism, genetic information processing, and environmental signal processing (Figure 3B). Furthermore, gene loss was observed across the fungal MAPK signaling pathway, with genes required for proper hyphal/cell wall growth absent from OAc genome, including *WSC1, RHO1, YPD1* and *SSN6*, which have all been implicated in hyphal generation and/or cell wall integrity (Verna, et al. 1997; Hwang, et al. 2003; Sohn, et al. 2003; Krysan, et al. 2005; Richthammer, et al. 2012).

**Figure 4.**
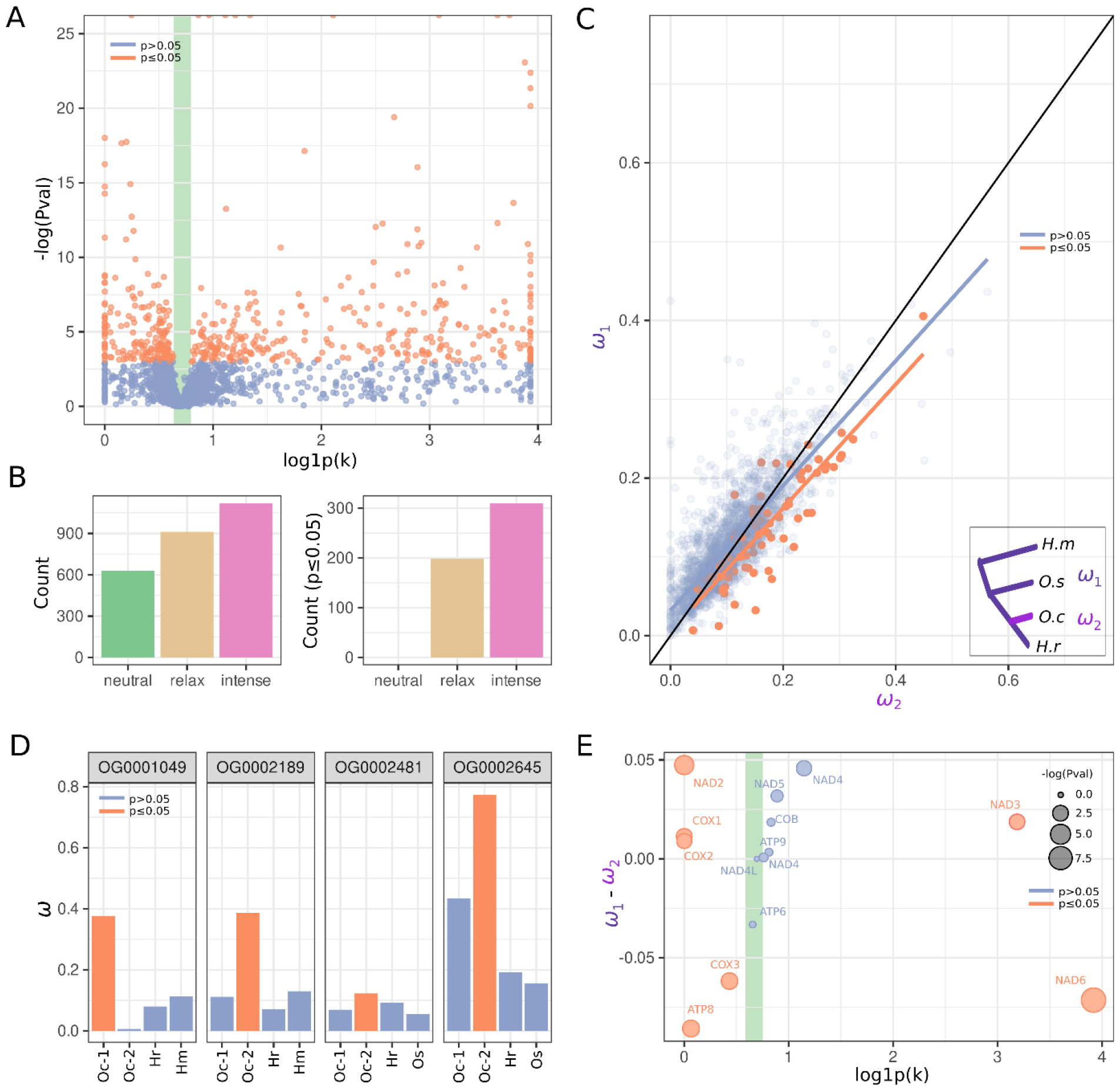
Widespread intensified and diversifying selection in the nuclear genome coupled with relaxed mitochondrial selection shapes the transition to symbiosis in OAc. A) Single copy ortholog (n = 2711) intensification parameter (*k*) and the significance of intensification (*k* ≥ 1.1)/relaxation (*k* ≤ 0.9) of selection observed in OAc against a background of *Ophiocordyceps sine, Hirsutella rhossiliensis* and *Hirsutella minnesotensis*. Green shading shows the expected value (0.9 < *k* < 1.1) under the null hypothesis that both lineages are being selected at similar rates. B) Left: the total number of single-copy orthologs with *k* ≥ 1.1 (intensified selection) and *k* ≤ 0.9 (relaxed selection) along the OAc terminal branch. Right: the number of single-copy orthologs with significant signal for intensification (*k* ≥ 1.1) or relaxation (*k* ≤ 0.9) of selection. C) dN/dS ratio (*ω*) calculated along the OAc terminal branch (*ω*_*2*_) and all other background branches (*ω*_*1*_) of the species topology shown in the tree. Single-copy orthologs were tested for diversifying positive selection to a p ≤ 0.05 cutoff. Solid colored lines represent the line of best fit calculated by glm for the significant and non-significant datapoints. Black line shows a linear relationship between *ω*_*1*_ and *ω*_*2*_. D) dN/dS ratio (*ω*) calculated across duplicated genes with positive selection. *ω* was calculated for all terminal branches of the tree shown in panel C and the significance level calculated for each terminal branch using a branch-site test for positive diversifying selection. E) Intensification parameter calculated across mitochondrial genes and the significance of the observed parameter. Green dashed line and shading shows the expected value (0.9 < *k* < 1.1) under the null hypothesis that both lineages are being selected at the near the same rate.

Toxic compounds, such as enterotoxins and mycotoxins, are an important mechanism for pathogenic *Ophiocordyceps* to kill or negatively impact their infected host (Lacadena, et al. 2007; Kobmoo, et al. 2018). To investigate if loss of toxin producing proteins had occurred during the transition to an associate life history in OAc, the ‘lost’ gene set was queried for genes involved in toxin production/pathogenicity. This identified 10 orthogroups related to toxin production that were absent from the OAc annotation, but which ranged from non-conserved (present in few taxa) to highly conserved (present in all taxa other than OAc) throughout *Ophiocordyceps* (Figure 3C). Two highly paralogous conserved orthogroups that were both predicted to encode Heat Labile Enterotoxins (HLEs) were found to be absent from OAc (Figure 3C; HLE). Four orthogroups that were predicted to encode proteins containing either Aflatoxin (Figure 3C; AfT) or Mycotoxin (Figure 3C; MyT) domains were also absent from the OAc lineage, along with other toxin domain containing orthogroups including Diphtheria-like toxin-like (Figure 3C; OG0000073) and ribotoxin (Figure 3C; OG0008875).

### Intensified and diversifying selection with potentially neofunctionalized paralogs shapes the transition to fungal symbiosis

Transition of life history from an entomopathogenic fungus to an associate likely requires complex changes in gene function and transcription. To understand the selection landscape of this transition, orthogroups were constructed using only species present in Clade 1 (Figure 3A) to maximize the number of nucleotide single-copy orthogroups for testing, identifying 2711 single-copy orthogroups for further analysis. Genome wide scans for relative changes in the intensity of selection in OAc against the background of non-associate *Ophiocordyceps* branches revealed 1120 (41.3%) orthologs that had *k* parameter values suggestive of intensified selection (*k* ≥ 1.1), in addition to 912 (34.4%) with values consistent with relaxed selection (*k* ≤ 0.9) (Figure 4A & B), while only 631 orthologs (23.3%) had a near-identical selective rate (1.1 ≥ *k* ≥ 0.9) across the entire phylogeny (Figure 4B). After testing for significant differences in evolutionary rate between the OAc and background non-associate *Ophiocordyceps* branches, 408 orthologs were identified that displayed a significant signal of either selection relaxation (n = 198) or intensification (n = 310) (Figure 4B), suggesting that intensified selective pressure, rather than relaxed selection, occurred more frequently in the OAc lineage since its separation from its most recent common ancestor (MCRA) with *H. rhossiliensis*.

To investigate the OAc genome for signatures of positive diversifying selection, *ω* (*d*_*N*_*/d*_*S*_) was calculated separately along the OAc terminal branch (Figure 4C; Foreground; *ω*_*2*_) and all other branches in Clade 1 (Figure 4C; Background; ω_1_) for each single-copy orthogroup. Comparing ω_1_ and ω_2_ values for each orthogroup revealed signals for elevated *ω* along the OAc terminal branch (*ω*_*2*_) compared to the background (*ω*_*1*_) (Figure 4C). Testing the foreground branch for diversifying positive selection in each single-copy ortholog identified 97 genes with significant signal of selection (p≤0.05) along the OAc lineage after its MCRA with *H. rhossiliensis* (Figure 4C). After filtering out orthologs with significant (p value ≤ 0.05) signal for selection across background branches, 82 genes were identified as displaying OAc specific diversifying. KEGG and GO functional analysis of these 82 genes under positive selection revealed functions involved in metabolism, cellular signalling and genetic information processing. With FAD binding (GO:0042254; p value = 0.019), ribosome biogenesis (GO:0042254; p value = 0.021), DNA-binding transcription factor activity (GO:0003700; p value = 0.025) and ATP-dependent chromatin remodeler activity (GO:0140658; p value = 0.027) GO terms enriched within the positive selected set compared to the entire OAc gene space.

In addition to the metabolism pathway selection above, significant (p≤0.05) signals for positive selection were detected specifically within acetoacetate metabolism (Sup Figure 5). The three genes (acetyl-CoA acetyltransferase, acetyl-Coenzyme A acyltransferase 1 and succinyl-CoA-3-oxaloacid CoA transferase), which are involved in catalyzing the reactions from acetyl – Coenzyme A (CoA) ↔ acetoacetyl – CoA ↔ acetoacetate in the presence of CoA, showed clear signals for positive selection that were absent from the background branches.

Twenty-three gene duplications were identified that were specific to the OAc lineage in the Clade 1 orthogroup set. Tests for diversifying selection on duplicated paralogs revealed four OAc duplications with significant (p value ≤ 0.05) signal. Duplicated paralogs had increased omega values compared to all other lineages within their respective gene trees (Figure 4D), suggesting they may be undergoing neofunctionalization through directional selection. Gene function of putatively neofunctionalized genes included the Zinc Finger Protein 598 (OG0001049), TPR-like (OG0002189), Fungal specific transcription factor (OG0002481) and DNA repair protein MutS (OG0002645) domain containing proteins.

In contrast to intensified selection across nuclear orthogroups (Figure 4A), mitochondrially-encoded genes exhibited frequent relaxed selection along the OAc lineage (Figure 4E). Six mitochondrial genes exhibited significant (p≤0.05) signal for relaxed selection (*NAD2, COX1, COX2, COX3* and *ATP8*) whereas only two (*NAD6* and *NAD3*) showed significant intensification. However, the majority (n = 7) of mitochondrial genes were found to lack significant signal for disparate selection intensities (Figure 4E).

## Discussion

Long-read genome assembly of small insects is difficult due to the reduced amount of total DNA within individuals. To extract sufficient DNA researchers generally pool multiple individuals from the same clutch to carry out sequencing and assembly (Consortium 2010; You, et al. 2013; Yang, et al. 2019). Although this has benefits thorough the ability to carry out size selection, variant reads and those spanning recombination breakpoints can confound assembly algorithms resulting in fragmented assemblies (You, et al. 2013). In this study, genome assembly was performed from a single individual of *Parthenolecanium corni* producing contiguous assemblies for the insect and its *Ophiocordyceps* fungal associate.

Scale insects have a diverse set of reproductive life histories ranging from parthenogenic to sexual. *Parthenolecanium corni* is thought to have a parthenogenic life history, although males have been observed to exist within populations (Gimingham 1934; Santas 1985) and sexual reproduction is suggested to occur. In this study, almost no heterozygous genetic variation was observed, suggesting that the sequenced genome represents a highly-homozygous diploid, with only a small number of minor structural variants differentiating the two haplotypes. This may be due to the complexities of non-mendelian chromosome assortment during reproduction in *Part. corni* (Gavrilov and Kuznetsova 2004), as *Part. corni* exhibits a Comstockiella chromosome system (Gavrilov and Kuznetsova 2004; Gavrilov 2007), whereby paternally inherited chromosomes are condensed into a heterochromatic body in somatic cells and then eliminated (Hodson, et al. 2023). This, coupled with a suggested founder’s effect (Ward, et al. 2023), may explain the limited genetic diversity present in the sequenced genome, rather than parthenogenesis. Chromosome number (1n) variability is high in scale insects (Coccidae), with various scale species estimated to have 5 – 18 chromosomes (Gavrilov 2007), compared to 4-32 chromosomes in mealybugs (Pseudococcidae) (Gavrilov 2007). High levels of structural rearrangements were observed between the chromosome level genome of the mealybug *Planococcus citri* and the *Part. corni* genome sequenced in this study, with low levels of macrosynteny between the two taxa. Taken together this suggests that the chromosome evolution of scale insects and mealybugs are exceptionally complex and more research is needed to more fully understand their evolutionary and population genetics.

Changes in gene function when transitioning life histories are well recorded, with strong selective pressures being imposed across the genome as the transitioning organism is exposed to novel stressors (Schluter, et al. 1991; Kelly and Griffiths 2021). Symbionts are no exception with rapid diversification of key mechanisms such as metabolism and growth being observed (Brownlie, et al. 2007; Wernegreen 2015). After adaptation to their host, relaxed selection is generally observed in the genome as the symbiont becomes shielded from many of the selective pressures that affect their free-living sister taxa (Wernegreen 2011; Wertheim, et al. 2015). Changes in filamentous vegetative growth, production of fruiting bodies and metabolism efficiency have all been suggested to be key pathways required for the transition to an internal associate life history. In the *Ophiocordyceps* associate of *Parthenolecanium corni* (OAc), strong intensification of selection was observed across the genome. Although the transition to an associate life history occurred before the species diversification of the *Parthenolecanium* genus (Szklarzewicz, et al. 2021), OAc may still be adapting to an internal, non-pathogenic life history and may not yet be providing symbiotic benefit to its host. Transition from a pathogenic life history may require more complex evolutionary changes to occur, as the associate must first prevent host death while rapidly switching to vertical inheritance. Despite this, pathogenic fungi already have the ability for intracellular survival during the initial infection stages of their life cycle (Kobmoo, et al. 2018), potentially requiring fewer alterations in cell growth and response to abiotic and biotic stimuli to occur.

Gene loss and genome shrinkage is a common occurrence during the transition to an associate or symbiotic lifestyle (Moran and Mira 2001; Fan, et al. 2015; Osvatic, et al. 2023). Gene loss was observed in the *Ophiocordyceps* associate across many conserved genes with important functions in metabolism, cell wall integrity and gene regulation. During the adaptation to a symbiotic lifestyle, the associate or symbiont coopts fundamental processes and nutrients from the host, with insect fungal associates being shown to aid in host sterol metabolism (Noda and Koizumi 2003; Body, et al. 2013; Oliver 2023). During the transition from pathogenic to an associate life histories it was expected that evolutionary changes would occur in genes involved in fundamental pathways such as toxicity, cell growth, and sexual reproduction. In line with these predictions, gene loss was observed across genes involved in hyphal growth and cell wall integrity as part of the fungal MAPK signaling pathway, suggesting that filamentous growth may not be necessary in this *Ophiocordyceps* associates. Therefore, the *Ophiocordyceps* associate of *Part. corni* may have reverted to a yeast-like life history as has been observed in other independent transitions in fungal insect associates (Suh, et al. 2001).

Pathogenicity and host control traits in *Ophiocordyceps* is suggested to be underpinned by complex interactions between toxin-related genes and the host organism (Kobmoo, et al. 2018; Will, Attardo, et al. 2023; Will, Beckerson, et al. 2023). Previous work has implicated Heat labile enterotoxins as compounds in *Ophiocordyceps* pathogenicity (Kobmoo, et al. 2018). Gene loss was also observed in multiple highly paralogous toxin genes including Heat liable enterotoxins which have been suggested to play an important role in *Ophiocordyceps* pathogenicity. Ribotoxins also play a key role in entomopathogenicity of Ophiocordyceptidae and other fungal taxa (Lacadena, et al. 2007; Foresman and Tartar 2023), which are thought to avoid exoskeleton drilling by favoring the production of ribotoxins early in the infection process (Foresman and Tartar 2023). However, only a single highly conserved ribotoxin gene was observed to be lost in the associate’s genome suggesting that widespread loss of this gene family has not occurred. This suggests that key genetic pathways are only partially undermined through gene loss to reduce pathogenicity during the transition to an associate life history.

Diversifying selection provides an evolutionary mechanism for neofunctionalization and changes in the efficiency of preexisting genes for adaptive novelty (Brownlie, et al. 2007; Murrell, et al. 2012). The *Ophiocordyceps* associate of *Part. corni* has undergone diversifying selection in multiple important mechanisms, such as genetic information processing, potentially changing the transcriptional landscape. In addition to changes in transcriptional control we observed diversifying selection across three sequential enzymes in the acetoacetate metabolism pathway suggesting that adaptive change has occurred across this pathway. Although this may be due to symbiotic production of extracellular acetoacetate for uptake by *Part. corni*, the presence of the *Ophiocordyceps* within the lipid bodies of *Part. corni* provides an attractive hypothesis (Szklarzewicz, et al. 2021). The *Ophiodordyceps* associate may be coopting nutrients from its host through metabolism of extracellular acetoacetate produced by *Part. corni*. This is supported by diversifying positive selection being observed in adjacent enzymes catalyzing the reversible reaction to acetyl-CoA. Furthermore, relaxed selection was observed within the mitochondrial genome providing support that the *Ophiocordyceps* associate may have lowered energy metabolism requirements than their pathogenic ancestors. Future work should investigate the validity of this hypothesis and determine the fitness of *Part. corni* without the associate as our results suggest that OAc may not be acting in symbiosis with its host.

This study investigated the evolutionary landscape during the transition from pathogenic to non-pathogenic associate life history in the *Ophiocordyceps* associate of *Parthenolecanium corni*. The results highlight metabolism and transcription as important mechanisms to reprogram during the transition. To achieve this, a long-read genome assembly was produced for *Parthenolecanium corni*, representing the first Coccoidea species assembled from a single individual, revealing divergent transposable element frequencies compared to other Hemipterans. Contiguous assembly of *Part. corni* allowed co-recovery of a chromosome level assembly for its *Ophiocordyceps* associate producing the first genome sequence for a non-pathogenic *Ophiocordyceps*. Investigation into the evolutionary history of the transition from pathogenic to associate life histories identified widespread gene loss of conserved orthogroups involved in toxicity, pathogenicity and transcriptional regulation. The selective landscape of the nuclear genome was intensified since the seperation of OAc from its closest pathogenic relative, contrasted with relaxed mitochondrial selection. Furthermore, we observed diversifying selection specific to OAc in genetic information processing and metabolism. Taken together our results suggest gene loss, transcriptional reprogramming and acetoacetate metabolism are important mechanisms for the transition from pathogenic to associate life histories in this *Ophiocordyceps*.

## Data availability

All associated read data can be found within NCBI BioProject PRJNA1083313. Gene annotations and their corresponding assemblies are deposited at 10.5281/zenodo.10780696.

## Acknowledgements

This work was funded by the Australian Wine Research Institute (AWRI) which is supported by Australian grape growers and winemakers through their investment body Wine Australia, with matching funds from the Australian Government. The AWRI is a member of the Wine Innovation Cluster (WIC) in Adelaide, Australia. Special thanks to grape growers that provided access to their vineyards and/or assisted with sample collection.

